## Supplementary material for "Recurrent connections facilitate occluded object recognition by explaining-away"

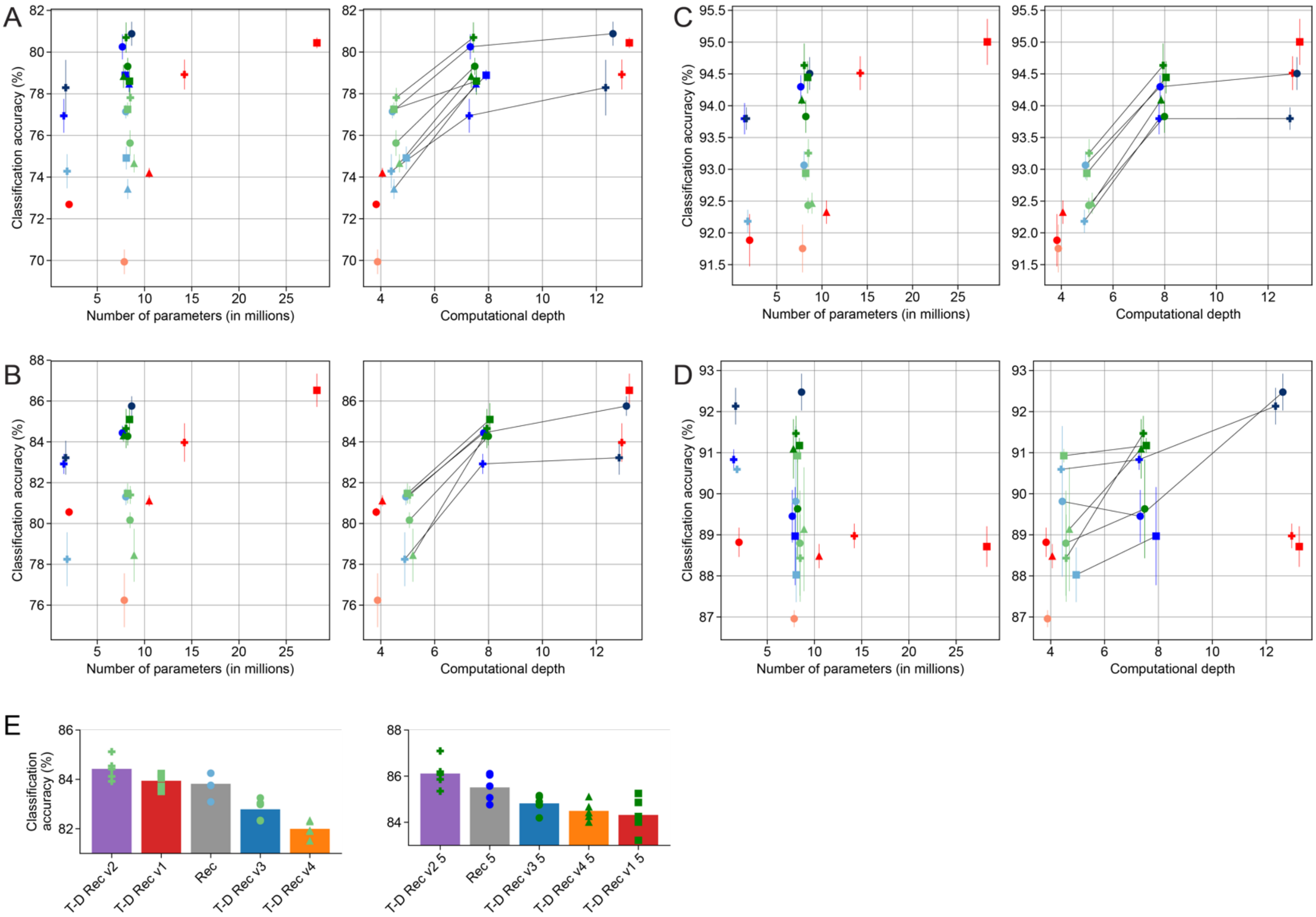


**Fig. S1.** Comparison of performance in the background-only, foreground-only, and unoccluded task. (*A*) Full task total accuracy vs. number of parameters and vs. computational depth. The total accuracy refers to the accuracy of predicting both the foreground and background object correctly. In case computational depths are different for the foreground and background prediction (e.g. as in the recurrent models that predict the two objects sequentially), we averaged them. n=5 independent initializations for each model. We added a random Gaussian noise of zero mean and 0.5 (for the number-of-parameters plot) or 0.15 (for the computational-depth plot) standard deviation to each dot's x-coordinate to separate out overlapping dots. The architecture represented by each symbol is the same as in Fig. 2. (*B*) Same plots as in (*A*), but for the background-only task accuracy. (*C*) Same plots as in (*A*), but for the foreground-only task accuracy. (*D*) Same plots as in (*A*), but for the unoccluded task accuracy (the accuracy of predicting both objects correctly). (E) Comparison of task performance of recurrent and top-down recurrent models. n=5 independent initializations for each model. (*Left*) models that run for two time steps. (*Right*) models that run for five time steps. One variant, T-D Rec v2, outperformed the recurrent models, whereas another variant, T-D Rec v1, outperformed them when run for two time steps but not when run for five time steps. The other two variants, T-D Rec v3 and v4 did not outperform the recurrent models for either choice of recurrent time steps. This demonstrates that performance of a top-down recurrent model is sensitive to the choice of the layers between which top-down feedback exists. In addition, it suggests that top-down feedback does not necessarily improve performance.

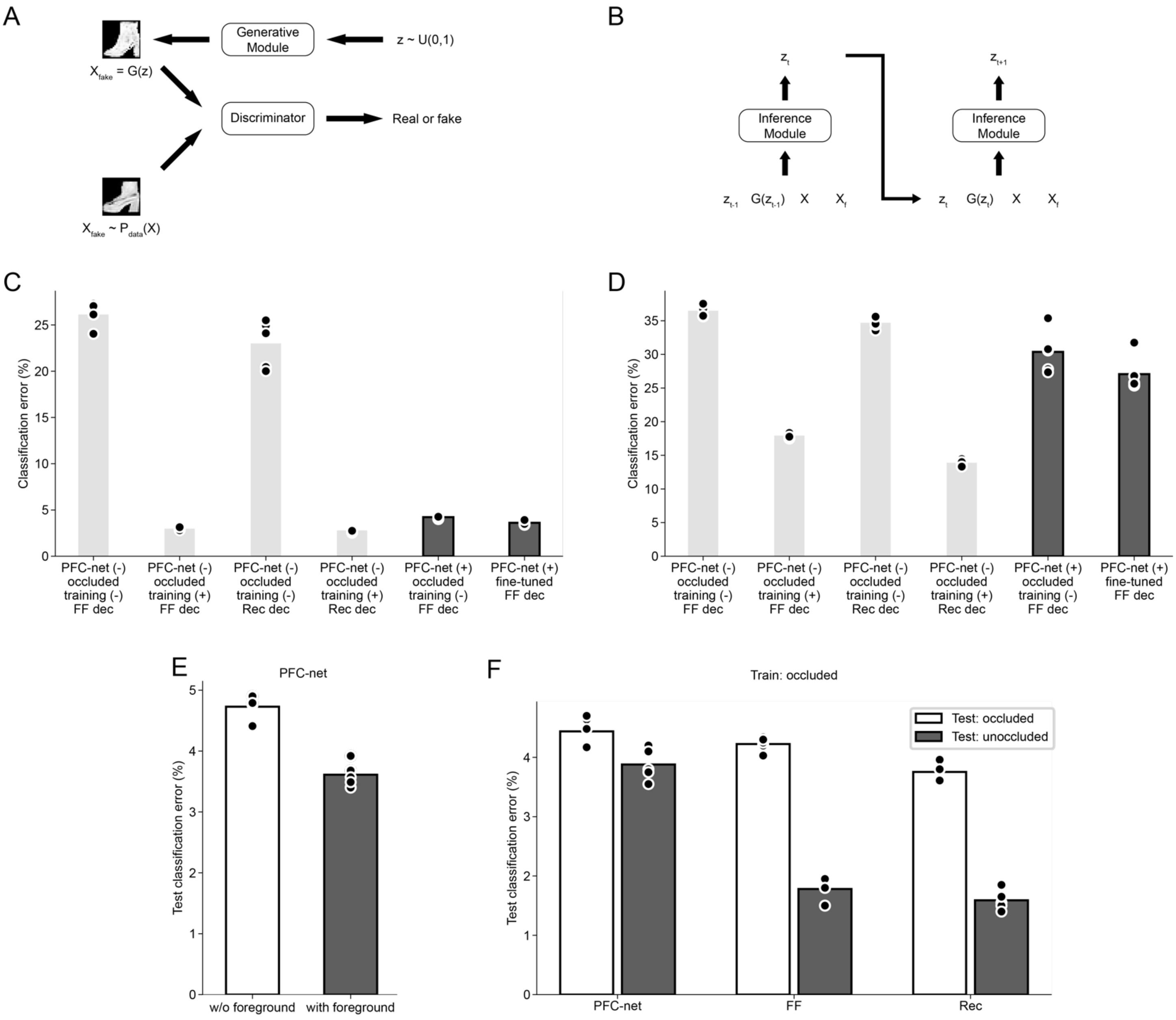


**Fig. S2.** Further details of PFC-net architecture and performance. (*A*) Schematics of generative module of PFC-net. (*B*) Schematics of inference module of PFC-net. (*C*) Same as Fig. 7*B*, but includes more models. As in Fig. 7*B*, "PFC-net (+ or -)" indicates whether the decoder takes PFC-net outputs or raw occluded images as input, and "occluded training (+ or -)" indicates whether the decoder is trained on occluded or unoccluded images. "fine-tuned" means that the decoder is fine-tuned on the outputs of PFC-net. "FF or Rec dec" indicates whether the decoder is feedforward or recurrent. n=5 independent initializations for each model. (*D*) Same as (*C*) for the predicted pose condition, where the spatial transformations to the background object is inferred and reversed by separately trained networks. (*E*) Comparison of PFC-net models with and without the foreground object image as an additional input (in the original pose condition). n=5 independent initializations for each model. (*F*) Occluded-to-unoccluded generalization. All three classes of models were trained on occluded images whose occlusion level ranged between 50% and 75% (see Methods), and tested either on held-out occluded images or unoccluded images. n=5 independent initializations for each model.

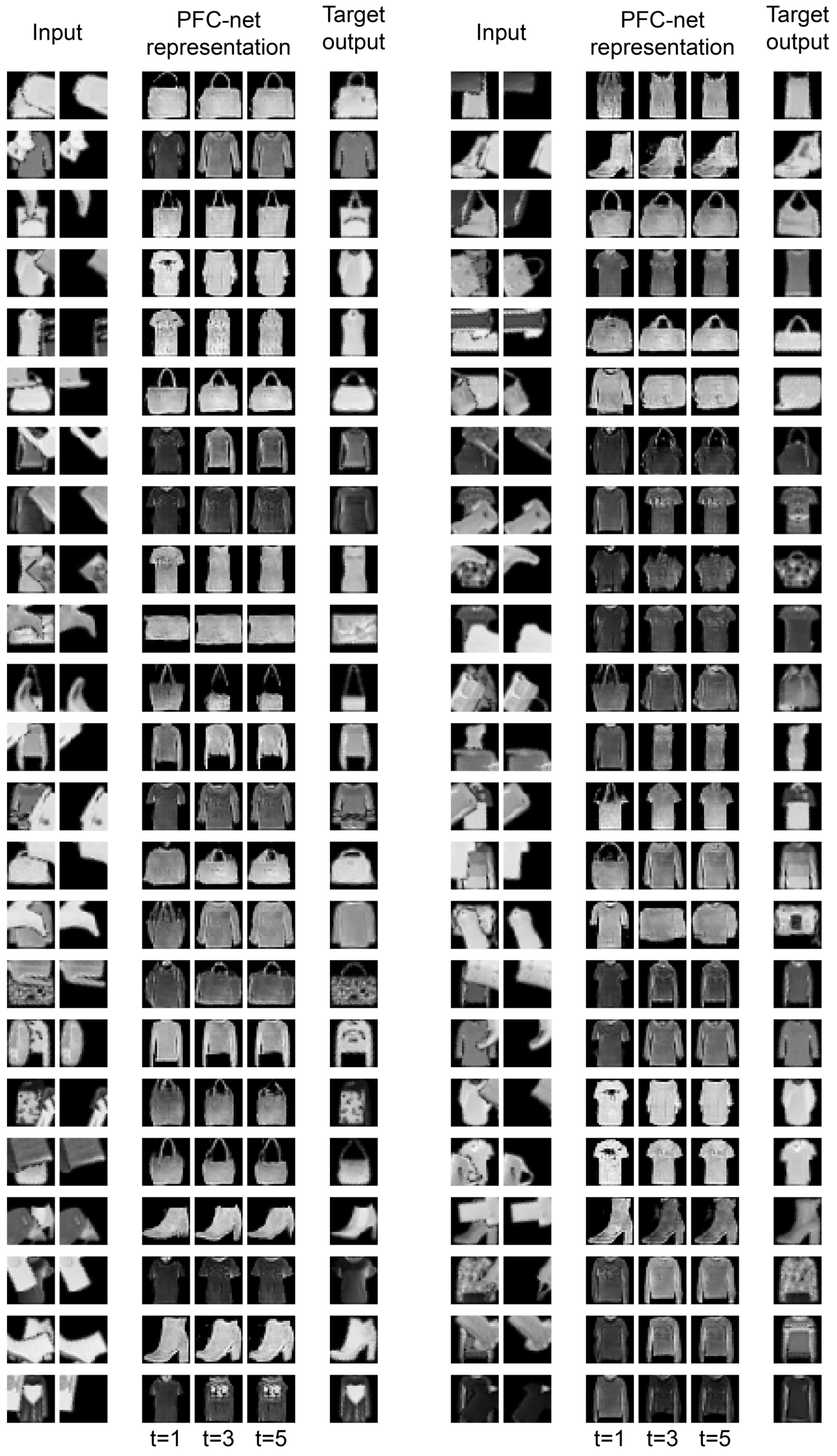


**Fig. S3.** More examples of PFC-net representation over time.

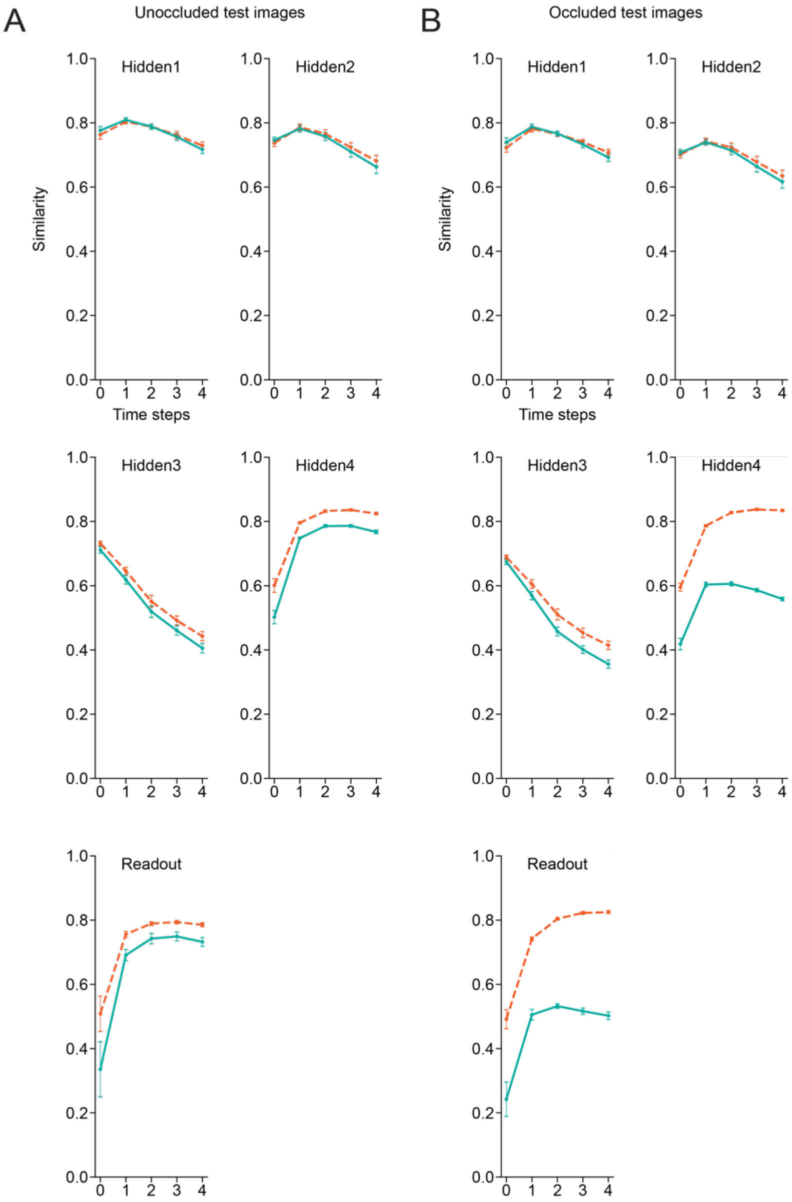


**Fig. S4.** Representational similarity between occlusion-trained and non-occlusion-trained recurrent networks in the random pose condition. Dotted orange lines indicate average representational similarity of occlusion-trained networks to other occlusion-trained networks across time steps. Solid green lines indicate average representational similarity of occlusion-trained networks to non-occlusion-trained networks. The error bars indicate the standard deviations across different pairs of randomly initialized instances of the networks (n=25 pairs for the solid green lines, and n=20 pairs for the dotted orange lines, excluding the pairs of the same instances). (*A*) When tested on unoccluded images. (*B*) When tested on occluded images.

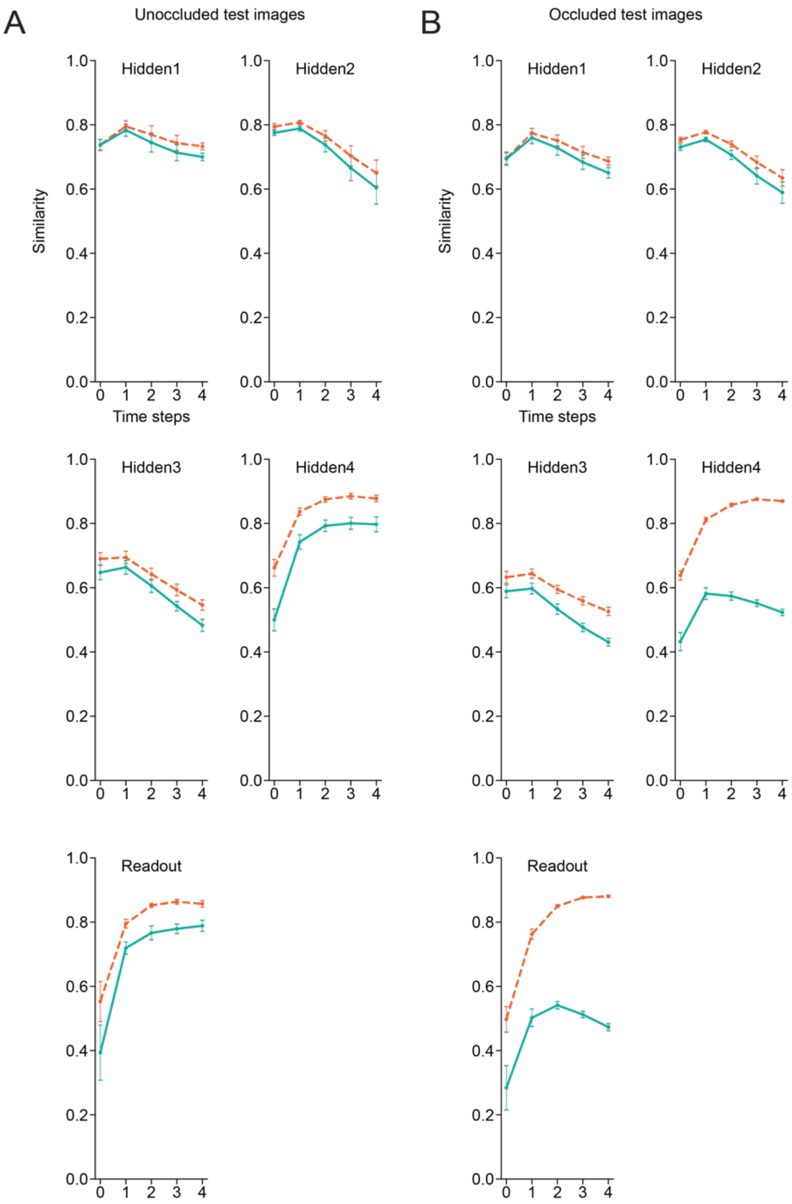


**Fig. S5.** Similar to Fig. S4 for top-down recurrent networks.

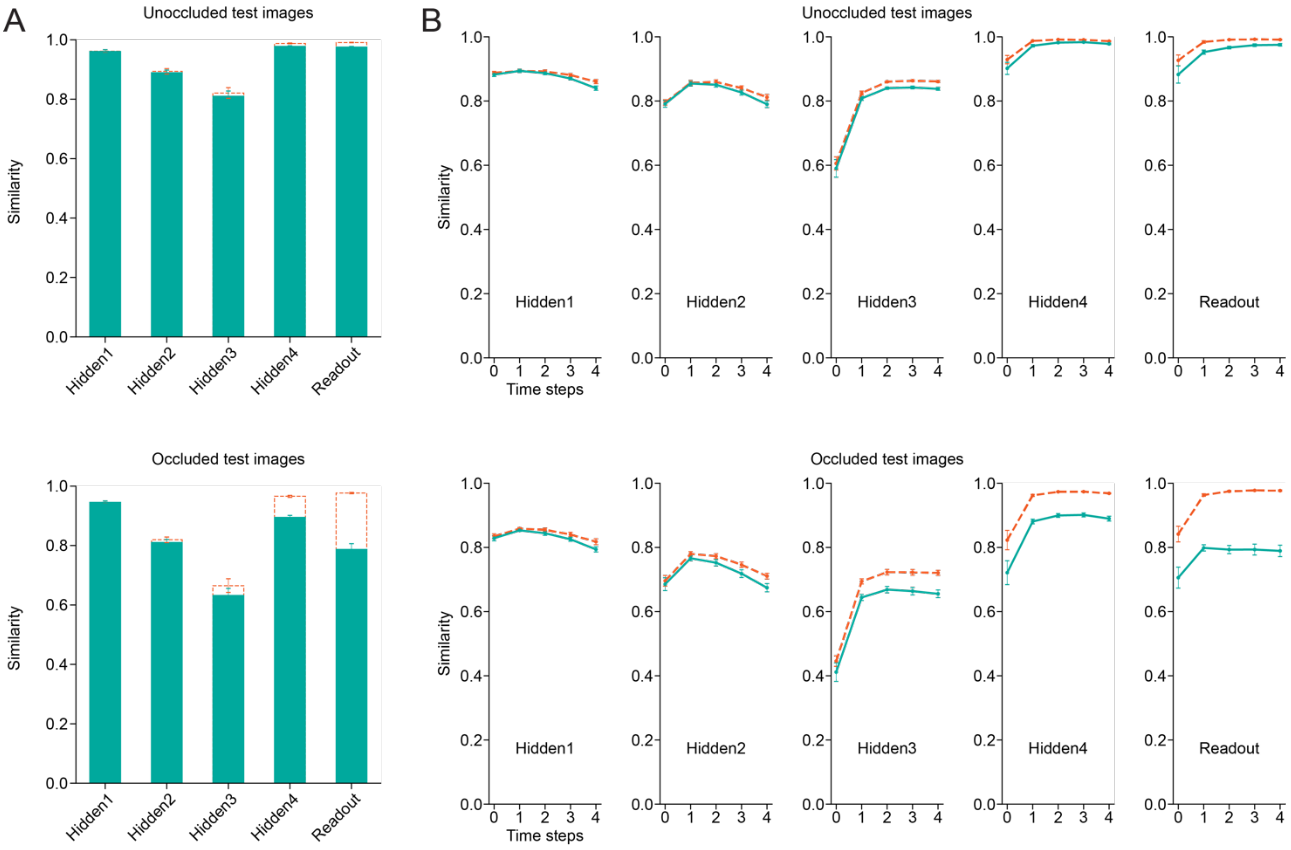


**Fig. S6.** Same as Fig. 8 for the original pose condition (with additional plots for the second and third hidden layers and readout layer of the recurrent architecture). (*A*) Representational similarity between occlusion-trained and non-occlusion-trained feedforward networks. (*Top*) when tested on unoccluded images. (*Bottom*) when tested on occluded images. (*B*) Same as (*A*) for recurrent networks.

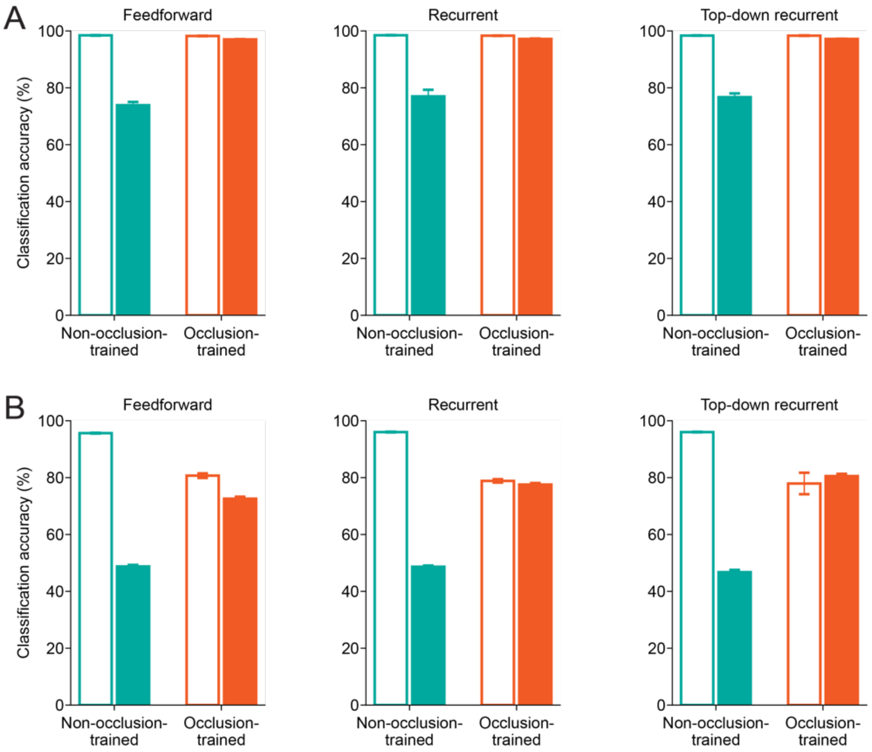


**Fig. S7.** Generalization performance of occlusion-trained and non-occlusion-trained networks. (*A*) Classification accuracy of occlusion-trained and non-occlusion-trained networks on occluded and unoccluded images in the original pose condition. In each bar graph, the green and orange bars represent non-occlusion-trained and occlusion-trained networks, respectively, and the empty and filled bars accuracy on unoccluded and occluded images, respectively. The error bars indicate the standard deviations across five independent initializations. (*Left*) feedforward architecture. (*Middle*) recurrent architecture. (*Right*) top-down recurrent architecture. (*B*) Same as (*A*) for the random pose condition.

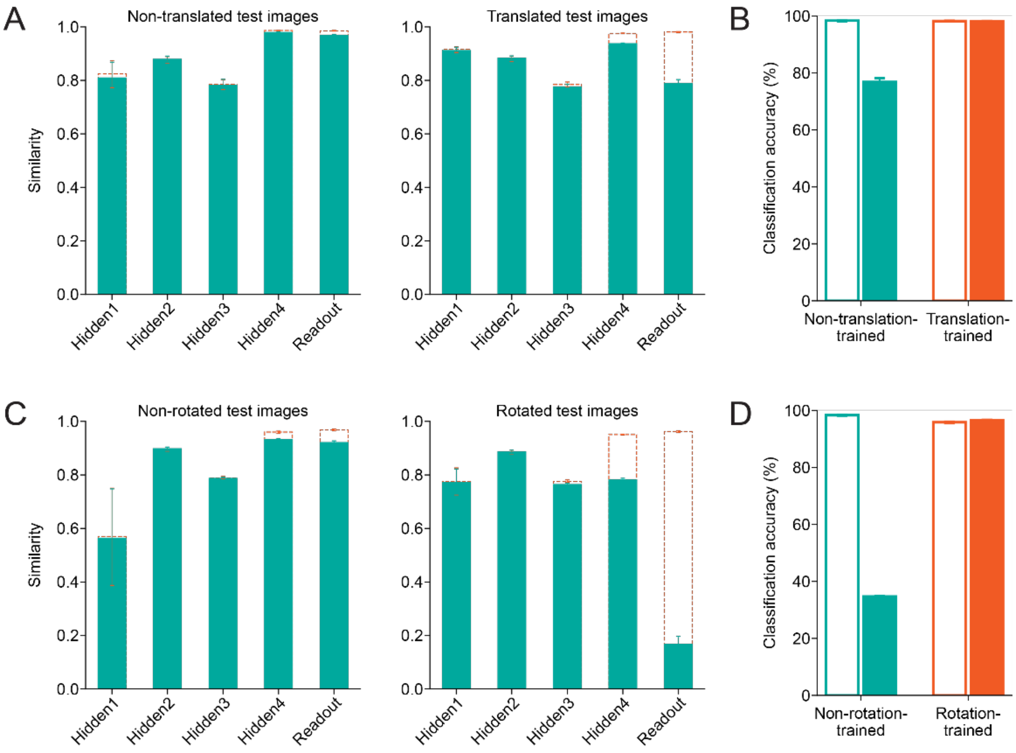


**Fig. S8.** Effect of spatial transformations on representational similarity. (*A*) Representational similarity between translation-trained and non-translation-trained feedforward networks. Dotted orange bars show average representational similarity of translation-trained networks to other translation-trained networks. Filled green bars show average representational similarity of translation-trained networks to non-translation-trained networks. The error bars indicate the standard deviations across different pairs of randomly initialized instances of the networks (n=25 pairs for the filled green bars, and n=20 pairs for the dotted orange bars, excluding the pairs of the same instances). (*Left*) when tested on non-translated images. (*Right*) when tested on translated images. (*B*) Classification accuracy of translation-trained and non-translation-trained networks on translated and non-translated images. The green and orange bars represent non-translation-trained and translation-trained networks, respectively, and the empty and filled bars accuracy on non-translated and translated images, respectively. The error bars indicate the standard deviations across five independent initializations. (*C*) Same as (*A*) for rotation. (*D*) Same as (*B*) for rotation.
